# A familiar face and person processing area in the human temporal pole

**DOI:** 10.1101/2023.10.15.562392

**Authors:** Ben Deen, Gazi Husain, Winrich A. Freiwald

**Affiliations:** Tulane University, New Orleans, LA, USA; The Rockefeller University, New York, NY, USA; Hunter College, City University of New York, New York, NY, USA

## Abstract

How does the brain process the faces of familiar people? Neuropsychological studies have argued for an area of the temporal pole (TP) linking faces with person identities, but magnetic susceptibility artifacts in this region have hampered its study with fMRI. Using data acquisition and analysis methods optimized to overcome this artifact, we identify a familiar face response in TP, reliably observed in individual brains. This area responds strongly to visual images of familiar faces over images of unfamiliar faces, objects, and scenes. However, TP did not just respond to images of faces, but also to a variety of high-level cognitive tasks that involve thinking about people, including semantic, episodic, and theory of mind tasks. The response profile of TP contrasted from a nearby region of perirhinal cortex that responded specifically to faces, but not to social cognition tasks. TP was functionally connected with a distributed network in association cortex associated with social cognition, while PR was functionally connected with face-preferring areas of ventral visual cortex. This work identifies a missing link in the human familiar face processing system that specifically processes familiar faces, and is well placed to integrate visual information about faces with higher-order conceptual information about other people. The results suggest that separate streams for person and face processing reach anterior temporal areas positioned at the top of the cortical hierarchy.

---

As we interact with familiar people such as friends, family and coworkers, their faces convey a wealth of information – from the essential question of who they are, to subtle cues about their feelings and thoughts. Behavioral research has demonstrated that familiar faces are perceived more effectively than unfamiliar faces, with more robust recognition across changes in viewpoint, expression, and context, leading to arguments for a qualitatively distinct system (*1-4*). What processes in the brain underlie the perception of familiar faces?

Models of familiar face perception have long argued for the presence of a “person identity” node, involved in connecting faces with social identity representations stored in long-term memory (Fig 1A, *5*). Neuropsychological evidence suggests that such a node might exist in the temporal pole (TP, also termed area 38 or TG), a primate-specific brain area at the anterior tip of temporal cortex. Patients with dysfunction of TP from brain damage, surgical resection, or frontotemporal dementia have been found to have difficulties recalling semantic information about familiar people, and/or connecting familiar faces with person identities (associative prosopagnosia, *6, 7, 8*).

**Figure 1:**
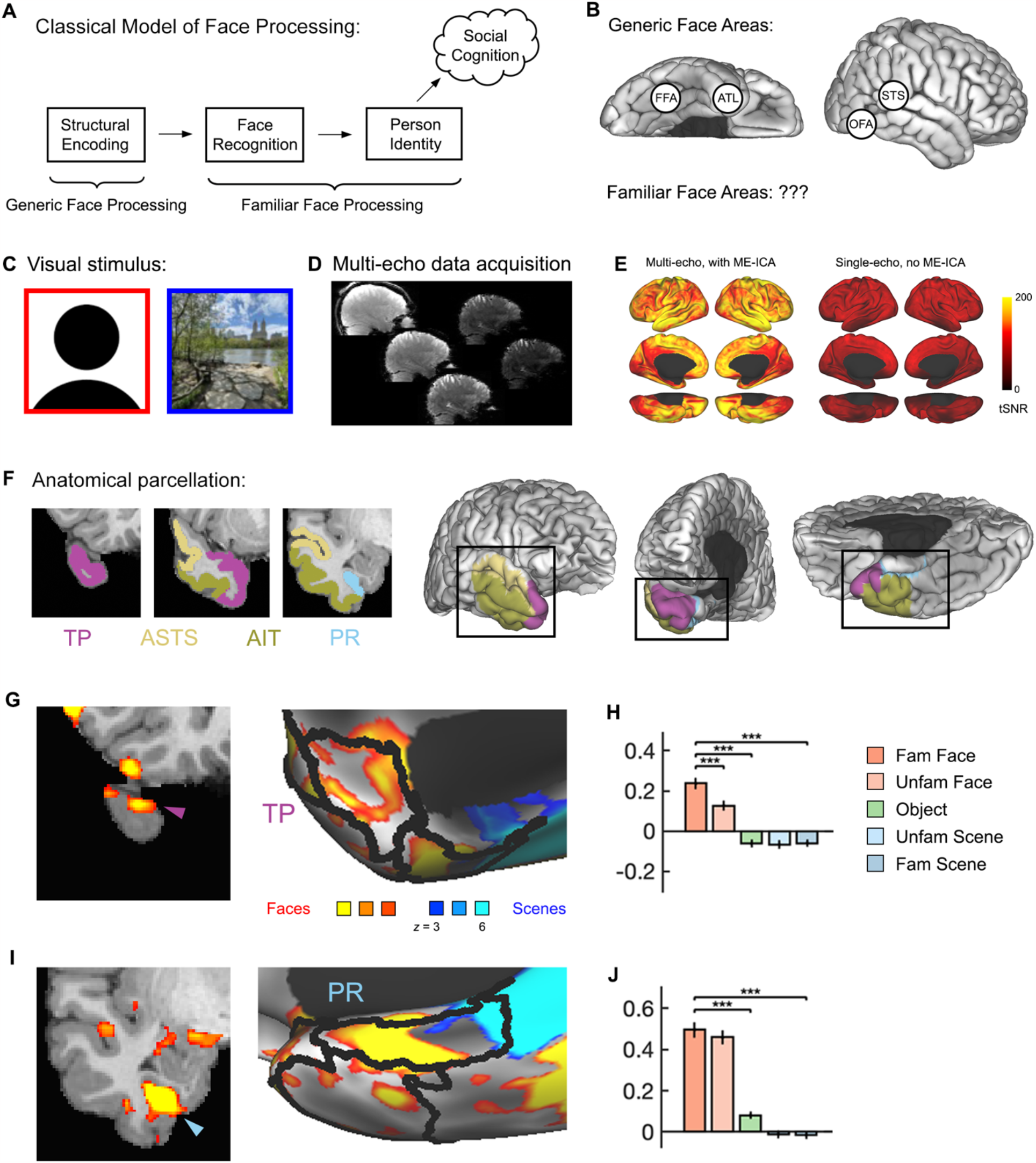
Face-preferring regions within the human temporal pole (TP). **A)** The classical model of cognitive operations supporting face recognition. **B)** Human brain areas that are known to respond to generic face images. **C)** Examples of face and scene images. **D)** Examples of raw data obtained using a multi-echo pulse sequence. **E)** Substantially higher tSNR was obtained using multi-echo data with multi-echo ICA preprocessing, relative to single-echo data with no ICA. **F)** Anatomical regions were hand-drawn to distinguish TP (area 38) from other neighboring regions of anterior temporal cortex: perirhinal cortex (PR), anterior superior temporal sulcus (ASTS), and anterior inferotemporal cortex (AIT). **G)** fMRI responses to faces versus scenes within TP (whole-brain general linear model-based analysis, thresholded at a False Discovery Rate of *q* < .01). **H)** Region-of-interest (ROI)-based responses across of face-preferring TP across visual conditions. * *P* < .05/4 = .0125, ** *P* < 10^−3^, *** *P* < 10^−4^ (ROI defined as the top 5% of face-preferring coordinates within anatomical search space; responses extracted from independent data; linear mixed model across runs, with participant included as random effect, error bars show standard error across runs). **I)** Responses to faces versus scenes within PR. **J)** ROI-based responses within PR.

However, neuroimaging studies of brain responses to familiar faces have not reliably found responses in TP (*9-13*). Studies presenting images or videos of unfamiliar and familiar faces have most commonly found responses in regions of posterior occipitotemporal cortex: the fusiform gyrus (fusiform face area, FFA), lateral occipital cortex (occipital face area, OFA), and posterior superior temporal sulcus (PSTS) (Fig 1B, *14, 15, 16*). A relatively weak response has been observed in ventral anterior temporal cortex, ventral and inferior to the temporal pole (anterior temporal lobe face area, ATL, *17, 18, 19*). Using images of personally familiar or famous faces, responses have additionally been observed in areas of association cortex – including medial prefrontal cortex (MPFC), medial parietal cortex (MPC), and the temporo-parietal junction (TPJ) – thought to be more broadly involved in social cognition (*11-13*). The lack of a reliable TP response to familiar faces may result from magnetic susceptibility artifacts in the anterior temporal lobes caused by the nearby presence of the sphenoid sinus, leading to lower temporal signal to noise ratio (tSNR) and power to detect effects in this area.

Our lab’s recent work has identified two face areas in anterior temporal cortex in the macaque: one in TP, and one in perirhinal cortex (PR, or areas 35-36, *20*). These zones of cortex have well-characterized cytoarchitecture and anatomical connectivity. Both areas have direct, bidirectional connections with the hippocampal formation via entorhinal cortex, and are thus well-positioned to support long-term memory for faces (*21-23*). The cytoarchitecture and anatomical organization of these regions are similar across humans and macaques (*24*). This raises the question: do humans have face-selective brain areas within the temporal pole and perirhinal cortex?

To address the question of whether humans have regions of TP and PR specialized for processing faces and/or people, we use three methodological advances. First, we present images of close personally familiar faces, to maximize the strength of mnemonic responses (Fig 1C).

Second, we use multi-echo pulse sequences optimized to maximize tSNR in the ventral anterior temporal lobes (Fig. 1D). We obtained substantially higher tSNR in the anterior temporal lobes using multi-echo data combined with multi-echo ICA preprocessing, relative to single-echo data without ICA (Fig. 1E). Third, in order to precisely characterize the anatomical organization of familiar face responses, areas TP, PR, AIT, and anterior superior temporal sulcus (ASTS) were hand-drawn on individual anatomical images based on previously established cytoarchitectonic and gross anatomical boundaries (Methods, Fig. 1D, *25, 26*). These methodological advances enabled the identification and precise anatomical characterization of anterior face responses in individual human brains.

Comparing responses to faces versus scenes – both familiar and unfamiliar – revealed face-preferring areas across multiple zones of the anterior temporal lobe (whole-brain general linear model-based analysis, corrected for temporal autocorrelation using prewhitening with an ARMA(1,1) model, and corrected for multiple comparisons across coordinates using a false discovery rate of *q* < .01; Figs. 1G, 1I, S2-3). Face-preferring responses were observed within TP in the left and right hemispheres of all participants (20/20 comparisons, Figs. 1G, 1I, S2A, S3A). Within a given brain and hemisphere, between one and three distinct regions were observed. The anatomical localization of these regions within TP varied, with responses commonly observed at the anterior-most tip of temporal cortex, in lateral aspects of the TP adjacent to ASTS, and in ventromedial aspects adjacent to the ventral anterior insula. By situating face responses within subject-specific anatomical parcellations, we were able to distinguish face responses in TP from nearby responses within ASTS, AIT, and PR. This result establishes the presence of reliable face responses within the human temporal pole, which can be identified within individual brains using optimized fMRI methods.

Face responses within PR were also observed across participants and hemispheres (20/20 comparisons, Figs. 1E-F, S2B, S3B). These regions were typically localized to the anterior-most aspect of the collateral sulcus, with some participants showing a second, more posterior region. This demonstrates the presence of a reliable face response within human perirhinal cortex, broadly consistent in anatomical location with the previously described ATL face patch. Based on this result, we propose that the ATL face area be more precisely named the PR face area. Brain regions within TP and PR preferring faces were similar in functional organization to face-responsive areas previously observed in the macaque (*20*). Given the similarity of cytoarchitectonic and anatomical organization of areas 38 and 35/36 across humans and macaques, this leads to the natural hypothesis that face-responsive regions of TP and PR are homologous across species. This contrasts from prior accounts, which have argued for a correspondence between anterior temporal face areas in humans with macaque area AM (*17*).

To what extent are neural responses within anterior temporal cortex selective for faces, among other categories of visual image? We addressed this question using a region-of-interest (ROI) analysis. Functional ROIs were defined as the top 5% of face-versus-scene-preferring coordinates within subject-specific anatomical search spaces, and response magnitudes across multiple categories were extracted in independent data. Throughout the remainder of the paper, the terms TP, PR, ASTS, and AIT will refer to face-preferring ROIs within these broader anatomical regions. We found that TP was strongly selective for faces over other categories (Fig. 1H). Responses to faces were above a resting baseline, while responses to scenes and objects were below baseline. Both left and right TP responded significantly more strongly to images of familiar faces than familiar scenes and generic objects (all *P*’s < 10^−8^, individual values in Tables S1-2; linear mixed effects model across runs, with participant included as a random effect).

Selective responses to faces were also observed in bilateral PR, ASTS, and AIT (familiar faces versus familiar scenes and generic objects, all *P*’s < 10^−4^, Figs. 1J, S4). This demonstrates that among visual stimuli, these functionally defined regions of anterior temporal cortex respond selectively to faces.

Neuropsychological results indicate a specific role for the temporal pole in processing familiar faces (*6, 8*). To what extent are TP responses modulated by familiarity? We compared responses to familiar and unfamiliar faces, using identities that were matched on age, race, and gender. Across both hemispheres, TP responded substantially and significantly more to familiar than unfamiliar faces (Fig. 1H, both *P*’s < .001, mean response increase 76%). In contrast, PR responded similarly to familiar and unfamiliar faces, with a weak but significant difference observed only in the left hemisphere (Fig 1J, left *P* < .0125 = .05/4, right *P* = .07, mean response increase 13%). This suggests a functional dissociation across anterior temporal face regions, in which PR contributes to processing all faces, while TP is specifically involved in processing familiar faces.

Is the functional role of TP limited to face processing, or does this area contribute more broadly to cognitive processes involved in understanding and remembering familiar people? To address this question, we scanned the same set of participants on an additional battery of tasks, eliciting visual, semantic, and episodic processing of other people, as well as theory of mind (ToM) reasoning. In the semantic task, participants rating personality traits of familiar people. In control conditions, they rated physical properties of generic objects, or spatial/navigational properties of familiar places. In the episodic task, participants imagined familiar people talking about specific conversation topics. In control conditions, they imagined generic objects engaged in physical interactions, or imagined navigating through familiar places. In the dynamic visual task, participants viewed images of unfamiliar faces, objects, and scenes (*27*). In the ToM task, participants read stories and answered true/false questions that either required understanding of a false belief or a false “photograph,” i.e. physical representation (*28*).

Consistent with the face preference described above, TP bilaterally showed an increased response to videos of unfamiliar faces versus objects and scenes (Fig. 2, dynamic visual task, all *P*’s < .005). However, TP also responded strongly to familiar people over object and place controls in the semantic task. Responses during trait judgment about familiar people were substantially and significantly stronger than responses while judging properties of familiar places and generic objects (all *P*’s < .005). TP also responded strongly to familiar person over object and place controls in the episodic task (all *P*’s < .0125). Lastly, right TP responded more when evaluating unfamiliar characters’ mental states in the ToM task, relative to physical states of the world (*P* < 10^−4^, left TP trending at *P* = .02). TP thus shows a strong and domain-specific response when processing familiar people across multiple tasks, which have a variety of stimulus features and behavioral demands. This suggests that TP is involved in social cognition and long-term memory – not just face perception – and is better characterized as person-selective than face-selective per se.

**Figure 2:**
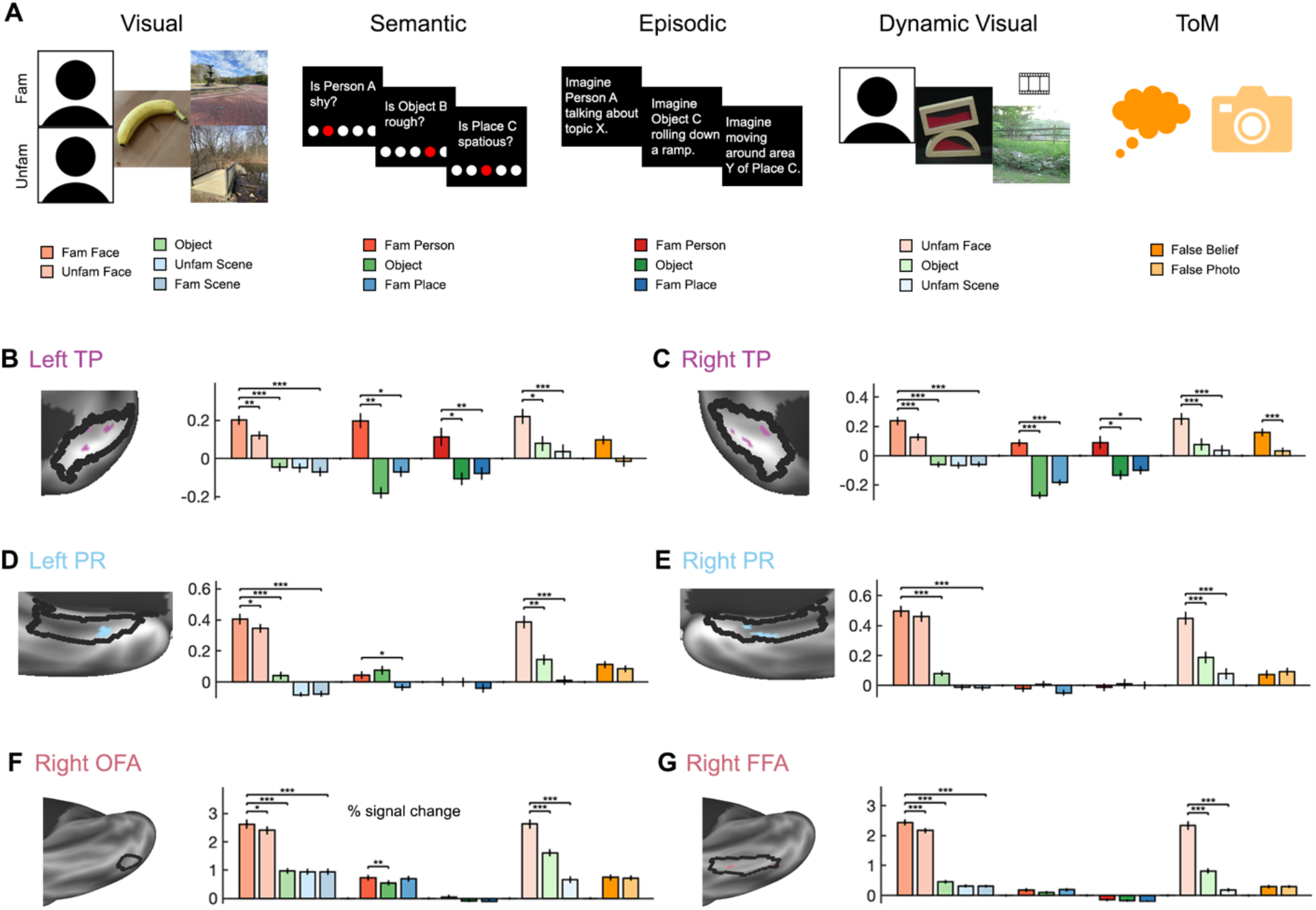
Region-of-interest (ROI)-based analysis reveals separate streams for person and face processing in anterior temporal cortex. **A)** Multiple tasks were used within individual participants, to determine whether areas responded during high-level social cognitive processing, in addition to face perception. Tasks included viewing images (visual task), trait judgment (semantic task), imagining events (episodic task), viewing videos (dynamic visual task), and reasoning about mental or physical representations (theory of mind task). **B)** ROI responses from the face-preferring part of left temporal pole (TP). **C)** Responses from right TP. **D)** Responses left perirhinal cortex (PR). **E)** Responses from right PR. **F)** Responses from the right occipital face area (OFA). **G)** Responses from the right fusiform face area (FFA). All ROIs were defined as the top 5% of face-preferring coordinates within anatomical search spaces. Responses (% signal change) were extracted from functionally defined ROIs across all task conditions. Error bars show standard error across runs. * *P* < .05/4 = .0125, ** *P* < 10^−3^, *** *P* < 10^−4^ (linear mixed model across runs, with participant included as random effect).

In contrast to the broad social responsiveness observed in TP, face-preferring regions of PR showed a strong response to faces but not other social conditions. PR responded bilaterally to videos of unfamiliar faces over scenes and objects (Fig. 2, all *P*’s < .001), but responded neither strongly nor reliably to people over places and objects in semantic and episodic tasks (left *P*’s ranging from .008 to 1; right *P*’s ranging from .09 to 1). A mixed effects two-way ANOVA across data from TP and PR identified an interaction between task condition and ROI (*P* < 10^−37^, *F*(17,3524) = 17.1) as well as main effects of condition (*P* < 10^−48^, *F*(17,3524) = 13.4) and ROI (*P* < 10^−11^, *F*(1,3524) = 43.1). Face-preferring regions of ASTS and AIT showed similar patterns of response to TP, with social preferences in both visual and cognitive tasks (Fig. S4). Response profiles thus contrast starkly among different face-preferring regions of anterior temporal cortex. This finding argues for two distinct streams of social information processing in the anterior temporal lobes: one for person processing, and another for face processing.

How are face-preferring areas of TP and PR situated as components of broader neural systems involved in social cognition and face perception? To ask this question, we defined face-preferring functional ROIs within anatomical search spaces covering brain areas previously implicated in social cognition and face processing. Regions of the social network included MPC, MPFC, TPJ, superior frontal gyrus (SFG), and middle superior temporal sulcus (MSTS). Regions of the face network included the FFA, OFA, and PSTS (Fig 3A).

**Figure 3:**
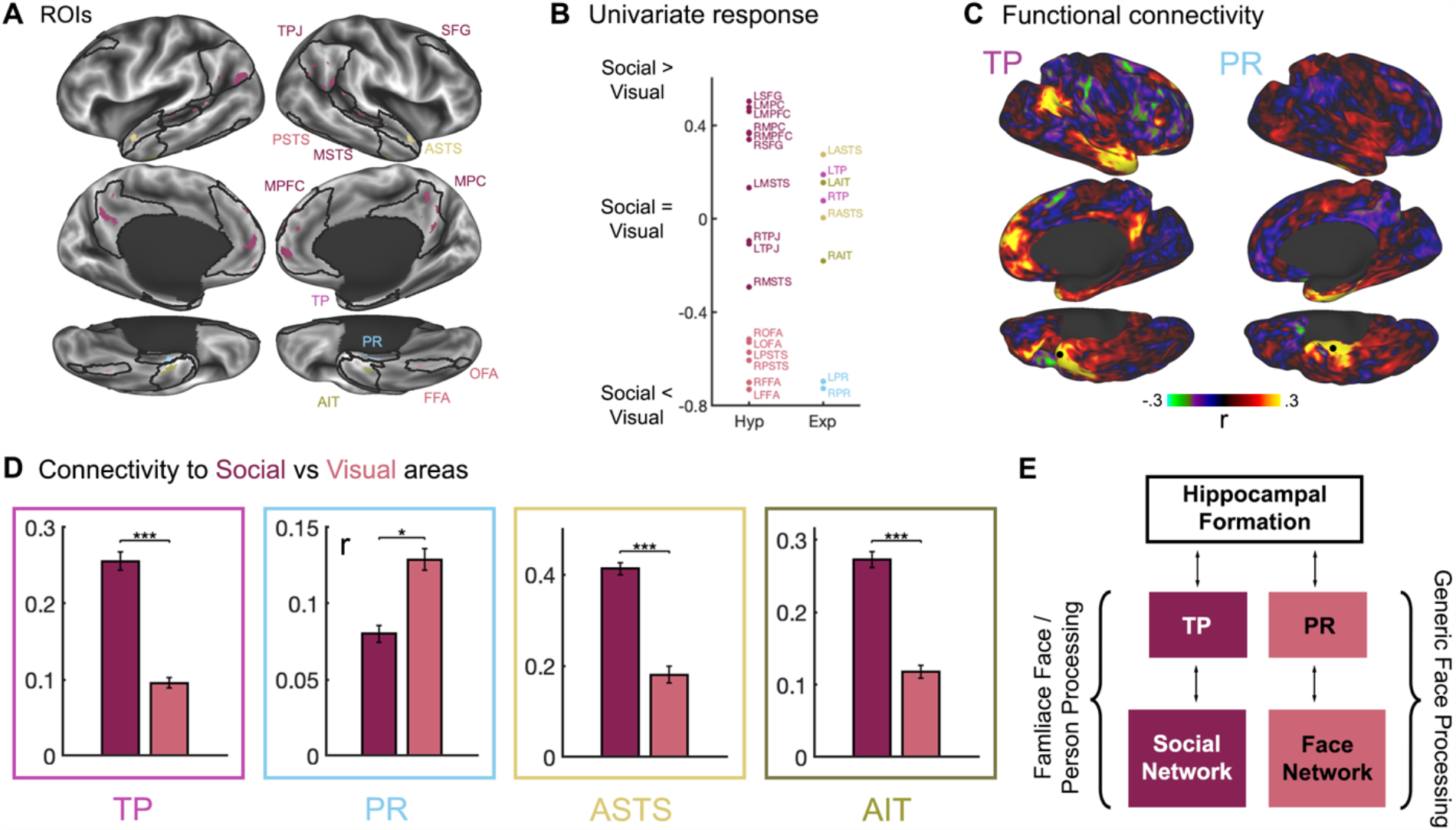
Functional connectivity shows that face-preferring temporal pole (TP) and perirhinal cortex (PR) are associated with separate large-scale networks. **A)** Functional ROIs shown in color, along with black outlines surrounding the anatomical search spaces used to define them. Social network regions include medial prefrontal cortex (MPFC), medial parietal cortex (MPC), superior frontal gyrus (SFG), temporo-parietal junction (TPJ), and medial superior temporal sulcus (MSTS). Face network regions include fusiform face area (FFA), occipital face area (OFA), and posterior superior temporal sulcus (PSTS). **B)** Plot of “social versus visual” modulation indices, capturing the relative degree of person selectivity in semantic and episodic tasks compared to the visual task. **C)** Whole-brain resting-state correlation maps from seeds in face-preferring right TP (left) and right PR (right). **D)** Correlations from face-preferring TP, PR, anterior superior temporal sulcus (ASTS) and anterior inferotemporal cortex (AIT) with areas of previously described social cognition and face perception networks. **E)** These results argue for separate streams for face and person processing within the anterior temporal lobe, providing separate bridges from the hippocampal formation to social cognition areas of association cortex, and to face perception areas in the ventral visual stream.

We first compare functional response profiles of TP and PR with social and face networks. Like TP, social network regions responded to people over places and objects across visual, semantic, and episodic tasks (all *P*’s < .0125, Fig. S5, Table S4-5). All social network regions except left MSTS also responded significantly more strongly to mental state versus physical reasoning in the ToM localizer (left MSTS *P* = .1, other *P*’s < .0125). In contrast, face network regions only responded strongly to visual face conditions over objects and scenes (all *P*’s < .0125, Fig. S6, Table S6). Among the other tasks, similar responses were observed to people, places, and objects, with only some small but significant differences observed in the semantic task (*P*’s ranging from .0002 to 1). Face network regions were not modulated by the ToM localizer, except for a small effect in right PSTS (right PSTS *P* < .001, other *P*’s > .2).

We computed a “social versus visual” modulation index, which captures the extent to which selectivity for people over object and place conditions is larger in cognitive tasks (social > visual), larger in the visual task (social < visual), or similar across cognitive and visual tasks (social = visual, Fig 3B). TP and most social network areas had stronger social selectivity in cognitive relative to visual tasks. In contrast, PR and face network areas had stronger modulation in visual relative to cognitive tasks. These results demonstrate that TP is similar in response profile to areas of the social network, while PR is similar to the face network.

To evaluate the functional network associations of TP and PR, we next measured synchronization of spontaneous fMRI signals in resting-state data. TP had reliable signal correlations with other social network regions within association cortex, including in MPFC, MPC, SFG, TPJ, and STS (Fig. 3C, S7). Face-preferring areas of TP – as well as ASTS and AIT – were significantly more correlated with social network areas than face network areas (Fig. 3D). In contrast, PR generally had low signal correlations with the rest of cortex, showing weak but positive correlations with the fusiform gyrus (Fig. 3C, S8), and significantly stronger correlations with face network versus social network areas (Fig. 3D). Taken together, these results indicate that TP constitutes a node in a broader network of association cortex involved in social cognition, while PR is instead related to areas of occipitotemporal cortex involved in face perception.

This study identifies a familiar face- and person-preferring region in the human temporal pole. These findings resolve a longstanding discrepancy between neuropsychological studies strongly indicating a role for the temporal pole in familiar face recognition and person understanding (*6-8*), and fMRI studies that have only rarely observed functional responses to familiar faces or people (*9-13, 29-31*). By presenting images of close personally familiar faces, and developing fMRI data acquisition and processing methods that boost signal quality in the anterior medial temporal lobes, we were able to reliably identify TP face responses in individual brains. By drawing anatomical regions based on established cytoarchitectonic criteria, face responses could be precisely localized to area 38. These results refine the view provided by neuropsychological studies, which typically involve alterations of large parts of cortex, by identifying focal brain regions with domain-specific responses.

Prior models of familiar face perception have posited that posterior-to-middle ventral temporal areas, such as the OFA and FFA, support an initial generic face processing step, while anterior temporal cortex supports subsequent person recognition and retrieval of high-level social information from long-term memory (*9, 16, 32, 33*). Contrasting with this view, we present evidence for distinct systems for generic face and familiar person processing within the anterior temporal lobe, in regions of TP and PR that have direct anatomical connections with the hippocampal formation. This argues for distinct processing streams for people and faces that extend to the top of the cortical hierarchy.

While prior fMRI studies have typically studied a specific cognitive or perceptual process and combined data across multiple participants in a standard template space, our approach assesses responses to a diverse set of tasks within individual human brains (*34-36*). This method enabled the discovery of a strong functional dissociation in the response profiles of TP and PR.

TP responded to a range of tasks involving familiar people, including semantic judgment and episodic simulation, while PR responded to images of faces but not social cognitive tasks. This dissociation fits well with prior neuropsychological results. Perirhinal cortex has been implicated in object memory and perception across humans, macaques, and rodents (*37, 38*). In contrast, damage localized to the human temporal pole typically doesn’t impair the recognition of unfamiliar objects or faces, but can impair familiar face recognition and person knowledge (*6-8, 10*). Thus, our results converge with neuropsychological evidence indicating separate streams for person and face processing within the anterior temporal lobe.

Face-preferring regions of TP and PR in humans had a similar functional organization to regions we have observed in macaques (Fig. 1G, *20*). The similarity in cytoarchitectonic and gross anatomical organization of areas 38 and 35/36 across humans and macaques (*24*) suggests that TP and PR face responses may be homologous across species. Further work comparing the connectivity and response profiles of these areas across species will be needed to evaluate this potential homology.

The functional connectivity of TP and PR indicate distinct interactions with large-scale brain networks, with TP signal correlated with a social cognition network distributed across association cortex, and PR more correlated with face processing areas in ventral temporal cortex. These differences fit well with anatomical connectivity patterns of ventral and temporopolar anterior temporal cortex established in the macaque. Cytoarchitectonic areas 38 and 35/36 both have anatomical connections with entorhinal cortex, but their connectivity with other parts of cortex differs. The temporal pole has strong connections with MPFC, orbitofrontal cortex, and dorsal STS (*23*), while perirhinal cortex shows strong connections with nearby inferotemporal cortex (*22*). PR is thus anatomically positioned as an interface between the hippocampus and the ventral visual stream, while TP is positioned as an interface between the hippocampus and transmodal association cortex.

The hypothesis of corresponding areas across species indicates that insight about the function of human anterior temporal cortex can be gained from invasive study of neuronal responses from these areas in macaques (*39, 40*). For instance, electrophysiological recordings from macaque TP have demonstrated that population responses encode the identity of familiar faces (*39*). These responses show a similar coding scheme to anterior parts of inferotemporal cortex – representing face identity in a way that is invariant to view angle – but only coding for personally familiar faces. In combination with the present results, this suggests that TP – while broadly involved in person understanding – nevertheless contains a visual representation of face features. On this hypothesis, TP is well positioned to support the integration of visual face processing with higher cognitive aspects of social understanding, such as the representation of mental state and trait information about familiar people.

## Methods

### Participants

Ten human participants (5 male, 5 female; age 28-40) were scanned using fMRI. Participants were healthy with normal or corrected vision, right-handed, and native English speakers. The experimental protocol was approved by the Rockefeller University Institutional Review Board, and participants provided written, informed consent.

### Tasks

Participants performed multiple tasks across three scan sessions. They were asked to choose six of their top ten most familiar people and places, and tasks involved processing these six people and places. Stimulus presentation scripts can be found at https://osf.io/5yjgh/.

In the main visual perception task, participants viewed serially presented naturalistic images of faces, objects, and scenes. For each participant, we obtained 20 images each of six familiar faces and scenes. Familiar face images were obtained directly from friends or family members, without the participant seeing them. Control images were defined as six yoked unfamiliar faces and scenes. Familiar and unfamiliar faces were matched on age group (young adult, middle-aged, old), race, and gender. Familiar and unfamiliar scenes were matched on rough semantic category (e.g. outdoor street view; building interior). Object images were of six generic objects with varying physical properties - a banana, a baseball, a feather, a rock, a sponge, and a wrench. Face and object images contained no clear spatial structure within the image (e.g. corners), and had minimal contextual cues beyond the background. Scene images contained no people. All five image categories were matched, for each participant, on the mean and variance across images of luminance, root-mean-square contrast, and saturation (in CIE-Lch space, with a D65 illuminant): all *P*’s > .05, one-way ANOVA and Bartlett test. Images were presented at 768 × 768 resolution, 12.8 × 12.8 degrees of visual angle, for 1.85s each with a 150ms interstimulus interval, in 18s blocks of images of a single identity. Participants performed a one-back task, pressing a button when an image was repeated. Task performance was high (mean hit rate 95.3%), indicating sustained attention throughout the scan. A post-scan questionnaire verified that participants could recognize the person or place in the familiar images (mean: 100% for faces, 84% for scenes), but not the unfamiliar images (mean: 0% for faces, 2% for scenes).

In order to evaluate the broader task sensitivity of face-preferring parts of cortex, four additional experiments were run. In the semantic task, participants rated traits of familiar people and places, and generic objects, on a 0 to 4 scale. This included personality traits of people (e.g. confident, angry, intelligent), spatial or navigational properties of places (e.g. cramped, large, has walls), and physical properties of objects (e.g. soft, heavy, rough). Participants rated by moving an icon left or right, over 18s blocks of four questions for a given identity, for a total of 20 questions per condition and identity.

Prior to the episodic task, participants listed five common conversation topics for each familiar person, and five familiar subregions of each familiar place. In the scan, they were asked to imagine familiar people talking about common topics, navigating specific subregions of familiar places, and objects engaged in physical interactions (e.g. rolling or sliding down a hill, falling into water). Participants were specifically asked to generate a novel event, rather than remember a past event. Imagination blocks lasted 18s, with a 3s verbal prompt, and a 1s hand icon at the end of the block, which participants responded to with a button press. After the scan, a free response description was collected, to ensure that participants could describe what they imagined. 9/10 participants were able to describe all events; the remaining one recalled 75% of events.

Across the visual, episodic, and semantic tasks, blocks were separated by 4s of resting fixation, and presented in five 8-13 minute runs per task, with palindromic block orders, counterbalanced across runs and participants. 18s fixation blocks were included in the beginning, middle, and end of the experiment to estimate a resting baseline.

Two localizer tasks from the existing literature were also run: a localizer for areas involved in theory of mind (ToM; reading and answering questions about stories involving false beliefs or false physical representations, *41*) and dynamic visual perception (watching videos of moving faces, objects, and scenes, *27*). Localizer tasks were split into 4-5 minute runs, with four runs for ToM and six for dynamic perception.

### MRI acquisition

Participants were scanned on a Siemens 3T Prisma across three 2.5-hour sessions, which included task acquisitions as well as 40 minutes of high-resolution anatomical images, 60 minutes of resting-state acquisitions, and spin echo acquisitions for distortion correction. Three each of T1- and T2-weighted anatomical images were acquired at .8mm resolution. Task and resting-state data were acquired using a multiband, multi-echo EPI pulse sequence, optimized to boost temporal signal-to-noise ratio (tSNR) in the anterior medial temporal lobes (TR = 2s, TE = 14.4, 33.9, 53.4, 72.9, and 92.4ms, 2.4x2.4x2.5mm resolution, 48 oblique axial slices with near whole brain coverage, multiband acceleration 3x, GRAPPA acceleration 2x, interleaved slice acquisition). 3-4 parameter-matched spin echo acquisitions were acquired per scan, between every four runs of task. Raw MRI data can be found at https://openneuro.org/datasets/ds003814. Separate analyses of this data have been presented elsewhere (*42*).

### MRI preprocessing

Data were preprocessed and analyzed using a custom pipeline, integrating software elements from multiple software packages: FSL (6.0.3), Freesurfer (7.1.1), AFNI, Connectome Workbench 1.5, tedana 0.0.10, and Multimodal Surface Matching (MSM). The code is available at https://github.com/bmdeen/fmriPermPipe/releases/tag/v2.0.2, with dataset-specific wrapper scripts at https://github.com/bmdeen/identAnalysis.

Anatomical images were preprocessed using an approach based on the Human Connectome Project (HCP) pipeline (*43*). The three images for each modality were linearly registered using FLIRT (*44*) and averaged; registered from T2- to T1-weighted; aligned to ACPC orientation using a rigid-body registration to MNI152NLin6Asym space; and bias-corrected using the sqrt(T1*T2) image (*45*). Cortical surface reconstructions and subcortical parcellations were generated using Freesurfer’s recon-all (*46*). Surface-based registration (MSMSulc) was used to register individual surfaces to fsLR average space (*47*). This registration was used to project the HCP multimodal cortical parcellation (*48*) onto individual surfaces.

Functional data were preprocessed using a pipeline tailored to multi-echo data, aiming to optimize tSNR while maintaining high spatial resolution. Motion parameters were first estimated using MCFLIRT (*49*). Intensity outliers were removed using AFNI’s 3dDespike, and slice timing correction was performed using FSL’s slicetimer. Motion correction was then applied, in combination with topup-based distortion correction (*50*), and rigid registration to a functional template image, in a single-shot transformation with one linear interpolation, to minimize spatial blurring. Multi-echo ICA was performed using tedana, with manual adjustments, to remove non-blood-oxygen-dependent (non-BOLD) noise components (*51*). Data were intensity normalized across participants. Data were subsequently analyzed in the volume (native functional space) and on the surface, after resampling to an individual-specific CIFTI space aligned with the anatomical template image, with 32k density fsLR surface coordinates, and 2mm volumetric subcortical coordinates. Registration between the functional and anatomical templates was computed using boundary-based registration (bbregister, *52*). Surface and volumetric data were both smoothed with a 2mm-FWHM Gaussian kernel. For resting-state data, the global mean signal was removed via linear regression, to diminish global respiratory artifacts not removed by multi-echo ICA (*53*). Pairs of time points with excessive head movement between them (framewise displacement > .5mm for task data, .25mm for resting-state data) were excluded from subsequent analysis. For the sake of comparing tSNR across single-and multi-echo datasets, single-echo data were defined by selecting the third echo, and preprocessed in the same way but without multi-echo ICA.

### Whole-brain analysis

Whole-brain statistical analyses in volumetric and CIFTI space were performed in individuals using AFNI’s 3dREMLfit, modeling temporal autocorrelation with a coordinate-wise ARMA(1,1) model (*54*). Results were thresholded using a false discovery rate of *q* < .01 (two-tailed) to correct for multiple comparisons. We compared responses to faces versus scenes in the main visual perception task (boxcar regressors convolved with canonical double gamma hemodynamic response function).

### Definition of anatomical regions

To precisely localize functional responses to anatomical regions, the left and right TP, PR, ASTS, and AIT were hand-drawn on individual brains. Regions were drawn on coronal slices of MNI-aligned images at 2mm resolution, using .8mm T1 images as a reference. They were subsequently resampled to 32k-vertex surface hemispheres, and then modified in the volume to remove any discontinuities observed on the surface.

Regions of TP and PR were defined based on previously established anatomical boundaries of cytoarchitectonic areas 38 and 35/36 (Fig 1D, *25, 26*). TP extended anteriorly to the tip of temporal cortex, and posteriorly to its border with PR. On its dorsomedial surface, the TP extended laterally to the lateral bank of the lateral-most temporopolar sulcus. On anterior-most slices in which the temporopolar sulcus was not visible, there was no lateral border. On its ventrolateral surface, the TP extended laterally to the medial lip of the inferior temporal sulcus, or if that wasn’t visible, the superior temporal sulcus. On the 3-4 anterior-most slices in which neither sulcus was visible, there was no lateral border. Anterior to the fronto-temporal junction (FTP, the first slice containing white matter connecting the frontal and temporal lobes, also termed the limen insulae), TP had no medial boundary. Posterior to the FTP, its medial boundary was the medial edge of the parahippocampal gyrus (PHG).

The border between TP and PR was placed at whichever of two landmarks was farther anterior: the anterior tip of the collateral sulcus, or one slice (2mm) anterior to the FTP. The posterior border was two slices (4mm) posterior to the last slices containing the head of the hippocampus, determined by the presence of the gyrus intralimbicus. The lateral border of PR varied from the midpoint between the fundus and lateral edge of the collateral sulcus, to the midpoint of the lateral edge of the collateral sulcus and medial edge of the occipitotemporal sulcus, depending on the depth of the collateral sulcus, based on previously established criteria (*25*). The medial border of PR in its first and last slices (2mm) was the medial edge of the PHG. In all other slices, the medial border of PR was positioned at the midpoint of the lateral bank of the collateral sulcus, defining the boundary between PR and entorhinal cortex.

ASTS and AIT were defined based on gross anatomical features, rather than previously described cytoarchitectonic boundaries. The posterior border of both areas was arbitrarily chosen as two slices (4mm) anterior to the posterior border of PR. This position was chosen so that the ASTS typically comprised roughly the anterior-most third of the main branch of the STS, not including parts of the STS after its posterior split and branching. The anterior border of each region was defined by the posterior borders of TP and PR. ASTS included cortex within the STS, extending to the dorsal and ventral lips of the sulcus. AIT included cortex between the ventral lip of the STS and the lateral edge of TP or PR.

In addition to TP, PR, ASTS, and AIT – the experimental focus of this study – we defined a set of anatomical areas intended to capture zones previously implicated in face perception and higher order social cognition. These areas were defined as combinations of parcels from the HCP multimodal parcellation (Table S3, *48*). Anatomical zones of the face network were defined by choosing regions of the multimodal parcellation corresponding to the location of previously described face responses in the FFA, OFA, and PSTS, using parcels defined by Julian et al. (2012) as a reference (*55*). Anatomical zones of the social network covered regions associated with the default mode network, including MPFC, MPC, SFG, TPJ, and MSTS. The hand-drawn ASTS ROI was subtracted from the MSTS ROI to avoid potential overlap.

### Region-of-interest (ROI) analysis

ROI-based analyses were conducted to assess how face-preferring brain areas respond across a range of task conditions. ROIs were defined as the top 5% of face-vs-scene-preferring CIFTI coordinates (main visual task) within search spaces corresponding the anatomical regions described above. To extract responses in the main visual task, a leave-one-run-out analysis was performed, in which ROIs were defined in all but one run of data, and responses extracted in the left out run. Statistics were performed on percent signal change values across runs, using a linear mixed effects model (MATLAB’s fitlme), with participant included as a random effect.

Responses to people were compared with places and objects, across visual, semantic, episodic, and dynamic tasks. Responses to familiar and unfamiliar faces were compared, as were responses to the belief versus false photo conditions of the ToM localizer. All comparisons were thresholded at *P* < .0125 < .05/4, applying Bonferroni correction across the four experimental ROIs TP, PR, ASTS, and AIT. Multiple comparison correction was not applied across left and right hemispheres, because separation by hemisphere was intended as an internal consistency check. Correction was also not applied across areas of the social and face networks, which are treated as previously described areas whose response profiles are tested only to confirm expected responses and compare to the experimental ROIs.

To evaluate differences between the response profiles of TP and PR, a condition-by-ROI mixed effects ANOVA was performed on data across all task conditions (MATLAB’s fitlme). Effects of condition, ROI, and intercept were included as random effects by participant. The condition-by-ROI interaction was not included as a random effect, because doing so led to model convergence issues.

### ROI-based functional connectivity analysis

Resting-state correlations were computed between the face-preferring ROIs defined above, including the experimental ROIs TP, PR, ASTS, and AIT, along with areas of the social and face networks. For each of the experimental ROIs, correlations to social versus face network areas were compared using a permutation test, permuting regions (10,000 iterations, two-tailed test, *P* < .0125 threshold). CIFTI-based whole-brain correlation maps, with functional ROIs as seeds, were computed for face-preferring regions of right TP and PR within each individual participant.

### Social versus visual response analysis

To compare regions based on the extent to which they respond specifically to person conditions, rather than just face images, we computed a “social versus visual” modulation index. The mean response (percent signal change) to people over objects and scenes (“person contrast”) was first computed for visual, semantic, and episodic tasks. The person contrast for the visual task was then subtracted from the mean person contrast across semantic and episodic tasks, and normalized by response range for each participant. This measure is positive when effects of social content are larger for semantic and episodic than for visual tasks.

## Data and code availability

Raw data are available at https://openneuro.org/datasets/ds003814. Stimulus materials are available at https://osf.io/5yjgh/. Analysis code is available at https://github.com/bmdeen/fmriPermPipe/releases/tag/v2.0.2 (generic analysis tools) and https://github.com/bmdeen/identAnalysis (dataset-specific wrapper scripts).

## Acknowledgements

We thank the staff at the Cornell Citigroup Biomedical Imaging Center for assistance with data acquisition, and Charles Lynch for helpful discussion on pulse sequence design. This work was supported by fellowships from the Helen Hay Whitney and Leon Levy foundations (B.D.), and the Center for Brains, Minds & Machines, funded by National Science Foundation STC award CCF-1231216 (W.A.F.).

## Author Contributions

B.D. and W.A.F. conceived the experiment. B.D., G.H., and W.A.F. designed the experiment. B.D. collected and analyzed the data. B.D. and W.A.F. wrote the paper.

