## Supplementary Info for "A familiar face and person processing area in the human temporal pole"

| | LTP | LPR | LASTS | LAIT |
|---|---|---|---|---|
| Face vs Scene (Visual) | 1.11E-09 | 7.01E-10 | 9.92E-07 | 8.57E-06 |
| Face vs Object (Visual) | 2.75E-11 | 4.62E-08 | 8.47E-08 | 2.54E-05 |
| Familiar vs Unfamiliar Face | 6.30E-04 | 2.66E-03 | 1.31E-05 | 4.00E-02 |
| Person vs Place (Semantic) | 2.72E-03 | 8.18E-03 | 9.00E-06 | 3.98E-05 |
| Person vs Object (Semantic) | 1.99E-04 | 1 | 1.21E-08 | 2.96E-03 |
| Person vs Place (Episodic) | 4.06E-04 | 3.64E-02 | 1.11E-07 | 2.37E-04 |
| Person vs Object (Episodic) | 5.32E-03 | 9.58E-01 | 1.52E-08 | 3.93E-03 |
| Face vs Scene (Dynamic) | 1.67E-10 | 1.07E-07 | 3.41E-03 | 1.34E-04 |
| Face vs Object (Dynamic) | 1.04E-03 | 5.65E-04 | 4.63E-05 | 2.44E-03 |
| Belief vs Photo | 2.42E-02 | 1.98E-01 | 3.69E-04 | 1.34E-01 |

**Table S1.** *P*-values from face-preferring ROI analysis in left anterior temporal regions. Linear mixed model across runs, with participant included as random effect.

| | RTP | RPR | RASTS | RAIT |
|---|---|---|---|---|
| Face vs Scene (Visual) | 1.20E-10 | 1.71E-10 | 1.30E-11 | 1.49E-12 |
| Face vs Object (Visual) | 5.33E-15 | 2.26E-09 | 2.22E-16 | 2.27E-09 |
| Familiar vs Unfamiliar Face | 2.16E-05 | 6.67E-02 | 4.21E-09 | 6.05E-09 |
| Person vs Place (Semantic) | 1.69E-10 | 9.27E-02 | 0 | 3.40E-02 |
| Person vs Object (Semantic) | 6.38E-10 | 1 | 1.33E-15 | 3.74E-02 |
| Person vs Place (Episodic) | 1.25E-03 | 1 | 1.31E-04 | 4.57E-03 |
| Person vs Object (Episodic) | 1.51E-03 | 1 | 8.28E-06 | 5.21E-02 |
| Face vs Scene (Dynamic) | 4.59E-11 | 2.28E-07 | 3.25E-04 | 7.97E-08 |
| Face vs Object (Dynamic) | 7.53E-08 | 8.84E-06 | 2.83E-05 | 1.93E-06 |
| Belief vs Photo | 4.24E-05 | 1 | 3.24E-09 | 3.90E-02 |

**Table S2.** *P*-values from face-preferring ROI analysis in right anterior temporal regions. Linear mixed model across runs, with participant included as random effect.

| **Search Space** | **Parcels** |
|---|---|
| *Social Network* | |
| MPC | RSC, POS1, v23ab, 7m, PCV, 31pv, 31pd, d23ab, 31a, 23d |
| MPFC | 8BM, a32pr, p24, d32, 9m, a24, p32, 10d, 25, s32, 10r, 10v |
| SFG | 9a, 9p, 8BL, 8Ad, 8Av |
| TPJ | PFm, PGi, PGs |
| MSTS | STSdp, STSvp |
| *Face Network* | |
| FFA | FFC |
| OFA | PIT |
| PSTS | TPOJ1 |

**Table S3.** How social and face network search spaces were defined in terms of anatomical parcels from the multimodal parcellation (Glasser et al. 2016).

| | LMPC | LMPFC | LSFG | LTPJ | LMSTS |
|---|---|---|---|---|---|
| Face vs Scene (Visual) | 3.18E-09 | 6.44E-14 | 4.08E-08 | 1.33E-14 | 9.74E-10 |
| Face vs Object (Visual) | 1.23E-13 | 0 | 8.98E-10 | 1.33E-15 | 2.86E-10 |
| Familiar vs Unfamiliar Face | 1.06E-09 | 2.68E-12 | 8.89E-05 | 4.82E-11 | 1.83E-03 |
| Person vs Place (Semantic) | 4.08E-11 | 0 | 1.28E-10 | 2.87E-06 | 1.15E-02 |
| Person vs Object (Semantic) | 2.32E-13 | 0 | 7.21E-12 | 4.10E-06 | 1.37E-05 |
| Person vs Place (Episodic) | 1.44E-11 | 8.58E-10 | 4.55E-08 | 1.19E-08 | 3.16E-07 |
| Person vs Object (Episodic) | 2.55E-12 | 3.18E-14 | 2.93E-10 | 7.82E-06 | 1.81E-10 |
| Face vs Scene (Dynamic) | 1.20E-06 | 3.19E-09 | 2.19E-04 | 4.09E-07 | 4.62E-04 |
| Face vs Object (Dynamic) | 4.08E-04 | 7.35E-10 | 1.10E-03 | 4.89E-05 | 3.34E-03 |
| Belief vs Photo | 1.35E-07 | 1.34E-05 | 1.34E-05 | 1.75E-06 | 9.68E-02 |

**Table S4.** *P*-values from face-preferring ROI analysis in left social cognition areas. Linear mixed model across runs, with participant included as random effect.

| | RMPC | RMPFC | RSFG | RTPJ | RMSTS |
|---|---|---|---|---|---|
| Face vs Scene (Visual) | 5.49E-12 | 4.10E-12 | 5.22E-12 | 7.18E-11 | 2.84E-10 |
| Face vs Object (Visual) | 2.00E-15 | 1.65E-12 | 9.77E-15 | 1.98E-14 | 5.39E-12 |
| Familiar vs Unfamiliar Face | 3.42E-08 | 7.23E-10 | 5.06E-05 | 4.97E-10 | 1.33E-11 |
| Person vs Place (Semantic) | 4.38E-13 | 2.22E-16 | 1.54E-10 | 4.73E-06 | 3.51E-07 |
| Person vs Object (Semantic) | 3.11E-14 | 0 | 4.36E-11 | 4.31E-07 | 1.97E-07 |
| Person vs Place (Episodic) | 7.32E-09 | 1.23E-08 | 7.95E-05 | 1.49E-03 | 1.60E-04 |
| Person vs Object (Episodic) | 1.54E-10 | 1.41E-09 | 1.37E-06 | 5.49E-05 | 6.15E-06 |
| Face vs Scene (Dynamic) | 6.80E-09 | 3.29E-09 | 2.40E-05 | 1.01E-06 | 1.38E-05 |
| Face vs Object (Dynamic) | 5.22E-10 | 2.11E-07 | 2.11E-08 | 2.41E-04 | 1.20E-05 |
| Belief vs Photo | 1.09E-07 | 5.65E-05 | 1.00E-03 | 2.73E-07 | 1.30E-05 |

**Table S5.** *P*-values from face-preferring ROI analysis in right social cognition regions. Linear mixed model across runs, with participant included as random effect.

| | LFFA | RFFA | LOFA | ROFA | LPSTS | RPSTS |
|---|---|---|---|---|---|---|
| Face vs Scene (Visual) | 0 | 0 | 1.62E-11 | 1.58E-07 | 6.77E-06 | 3.16E-12 |
| Face vs Object (Visual) | 3.71E-13 | 0 | 1.27E-11 | 1.87E-10 | 1.82E-03 | 7.86E-13 |
| Familiar vs Unfamiliar Face | 8.68E-05 | 3.27E-06 | 5.96E-03 | 2.00E-03 | 5.39E-02 | 6.03E-10 |
| Person vs Place (Semantic) | 1 | 1 | 1 | 5.86E-01 | 1 | 4.39E-03 |
| Person vs Object (Semantic) | 1 | 1.25E-02 | 5.84E-03 | 1.81E-04 | 6.26E-01 | 2.23E-03 |
| Person vs Place (Episodic) | 4.15E-01 | 3.10E-01 | 4.23E-02 | 6.06E-02 | 4.49E-02 | 2.50E-02 |
| Person vs Object (Episodic) | 1 | 7.55E-01 | 2.51E-01 | 5.02E-02 | 8.77E-02 | 3.03E-02 |
| Face vs Scene (Dynamic) | 1.07E-10 | 1.22E-10 | 0 | 6.91E-08 | 1.40E-05 | 2.11E-12 |
| Face vs Object (Dynamic) | 3.65E-06 | 6.79E-09 | 0 | 2.20E-06 | 6.49E-04 | 5.84E-13 |
| Belief vs Photo | 1 | 1 | 1 | 6.95E-01 | 2.50E-01 | 6.18E-04 |

**Table S6.** *P*-values from face-preferring ROI analysis in bilateral face perception regions. Linear mixed model across runs, with participant included as random effect.

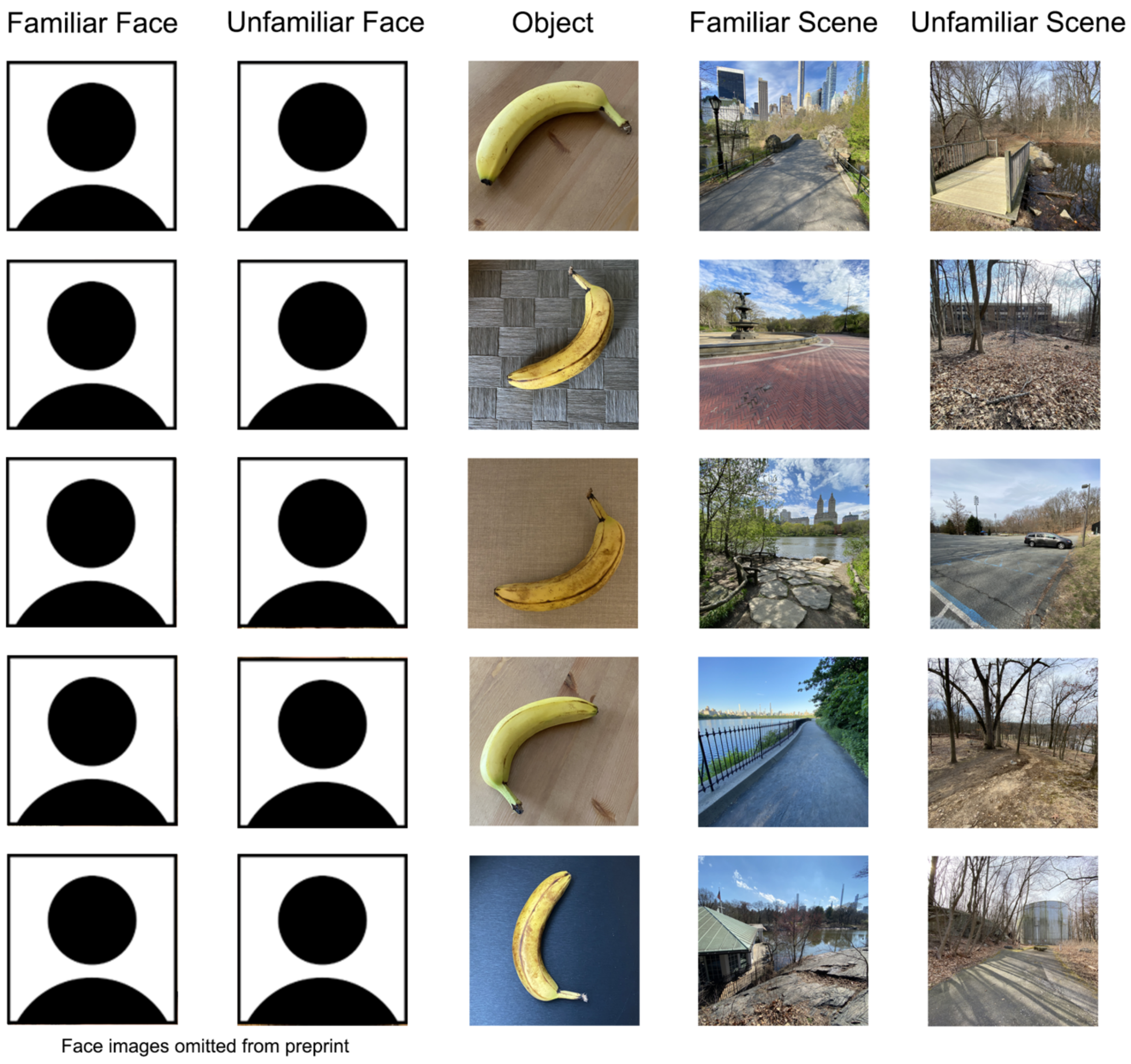

**Figure S1.** Additional examples of images used in the visual perception task. Each row contains examples from a given participant, condition, and identity.

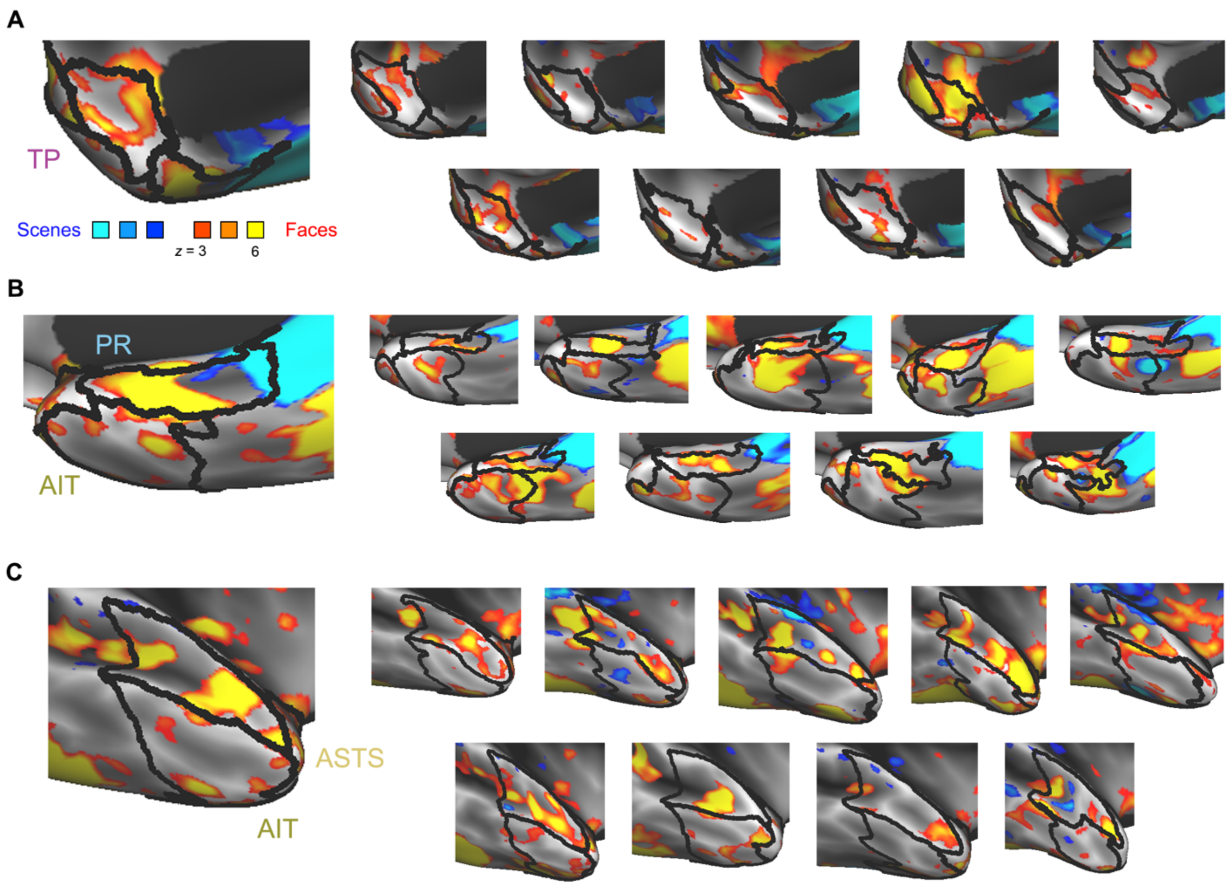

**Figure S2.** Individual participant surface-based responses to faces versus scenes within regions of right anterior temporal cortex: temporal pole (TP), perirhinal cortex (PR), anterior superior temporal sulcus (ASTS), and anterior inferotemporal cortex (AIT). Whole-brain general linear model-based analysis, thresholded at a False Discovery Rate of $q < .01$ to correct for multiple comparisons across coordinates.

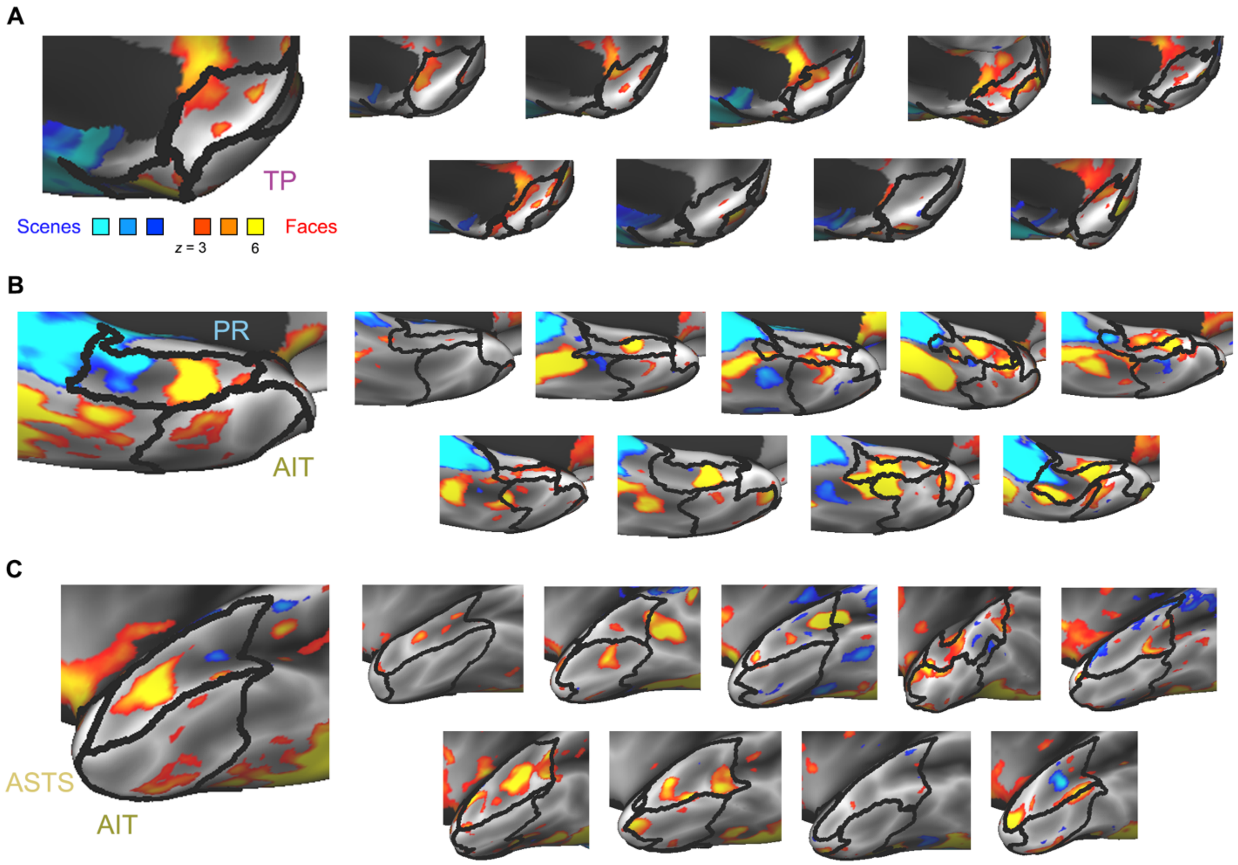

**Figure S3.** Individual participant surface-based responses to faces versus scenes within regions of left anterior temporal cortex: temporal pole (TP), perirhinal cortex (PR), anterior superior temporal sulcus (ASTS), and anterior inferotemporal cortex (AIT). Whole-brain general linear model-based analysis, thresholded at a False Discovery Rate of $q < .01$ to correct for multiple comparisons across coordinates.

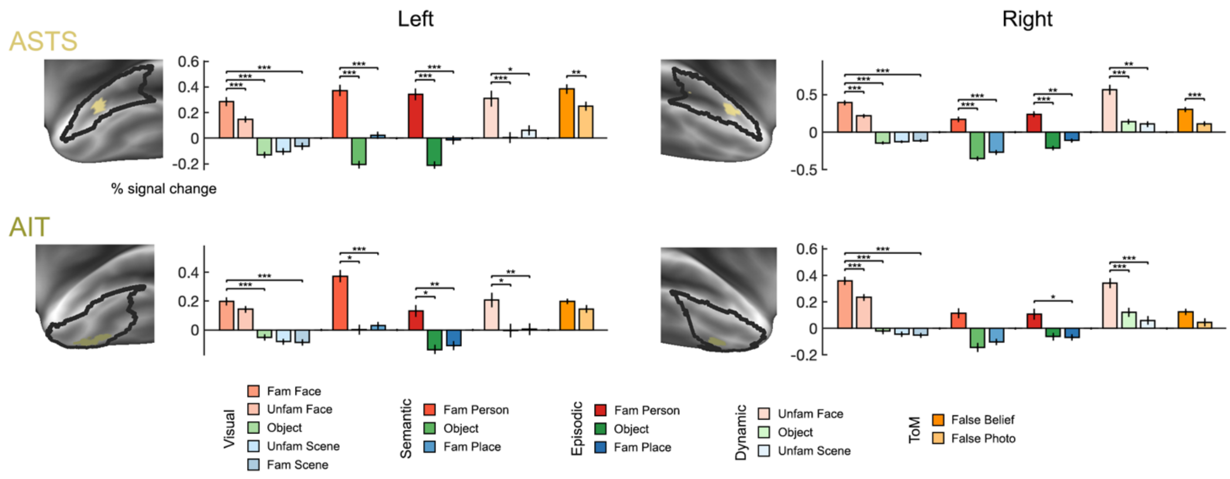

**Figure S4.** Region-of-interest (ROI) analysis in face-preferring regions of anterior superior temporal sulcus (ASTS) and anterior inferotemporal cortex (AIT). ROIs were defined as the top 5% of face-preferring coordinates within anatomical search spaces. Responses (% signal change) were extracted from functionally defined ROIs across all task conditions. Error bars show standard error across runs. * $P < .0125$, ** $P < 10^{-3}$, *** $P < 10^{-4}$ (linear mixed model across runs, with participant included as random effect).

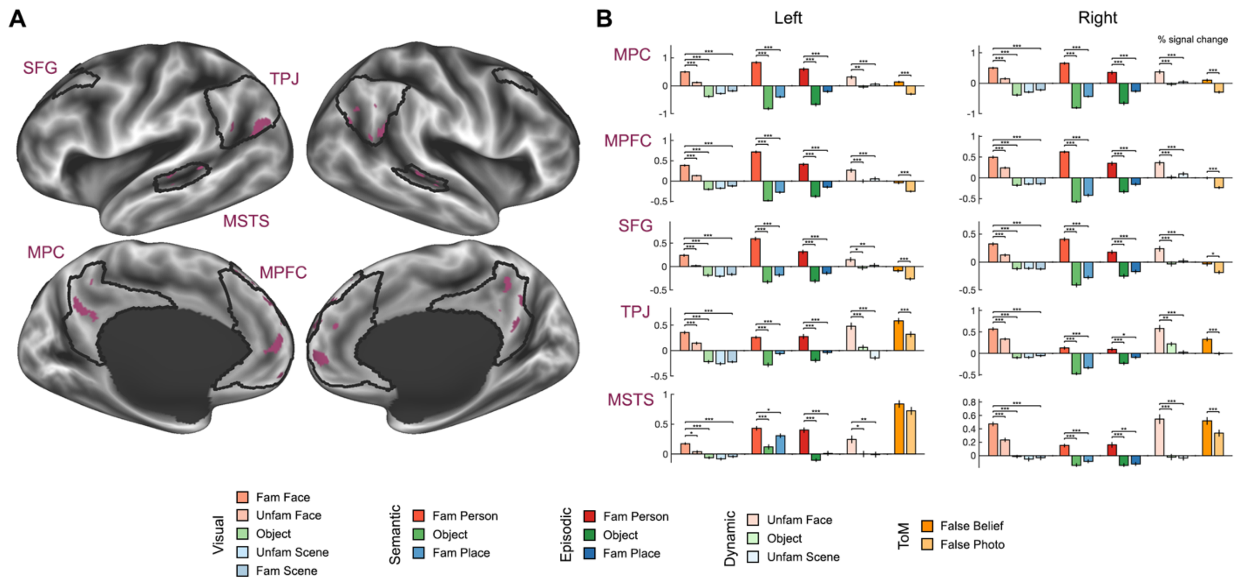

**Figure S5.** Region-of-interest (ROI) analysis in face-preferring regions within areas of association cortex implicated in social cognition. ROIs were defined as the top 5% of face-preferring coordinates within anatomical search spaces. Responses (% signal change) were extracted from functionally defined ROIs across all task conditions. Error bars show standard error across runs. * $P < .0125$, ** $P < 10^{-3}$, *** $P < 10^{-4}$ (linear mixed model across runs, with participant included as random effect).

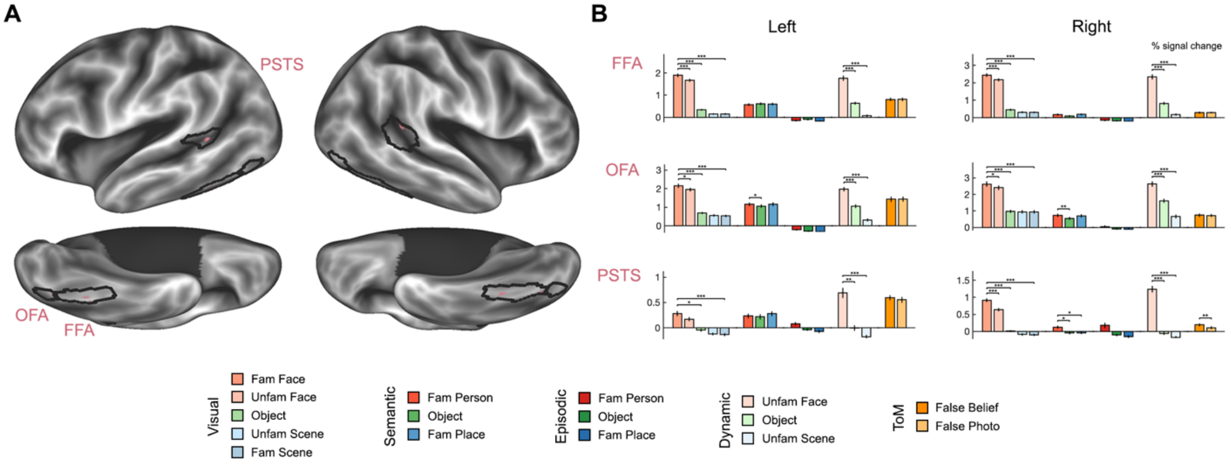

**Figure S6.** Region-of-interest (ROI) analysis in face-preferring regions within areas of cortex implicated in face perception. ROIs were defined as the top 5% of face-preferring coordinates within anatomical search spaces. Responses (% signal change) were extracted from functionally defined ROIs across all task conditions. Error bars show standard error across runs. * $P < .0125$, ** $P < 10^{-3}$, *** $P < 10^{-4}$ (linear mixed model across runs, with participant included as random effect).

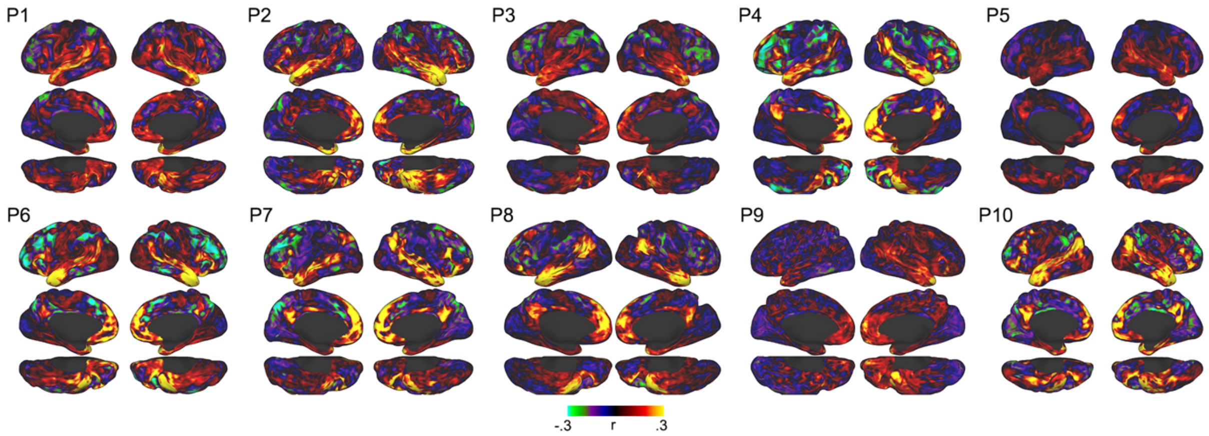

**Figure S7.** Whole-brain resting-state correlation maps from a face-preferring right temporal pole seed region, across all participants.

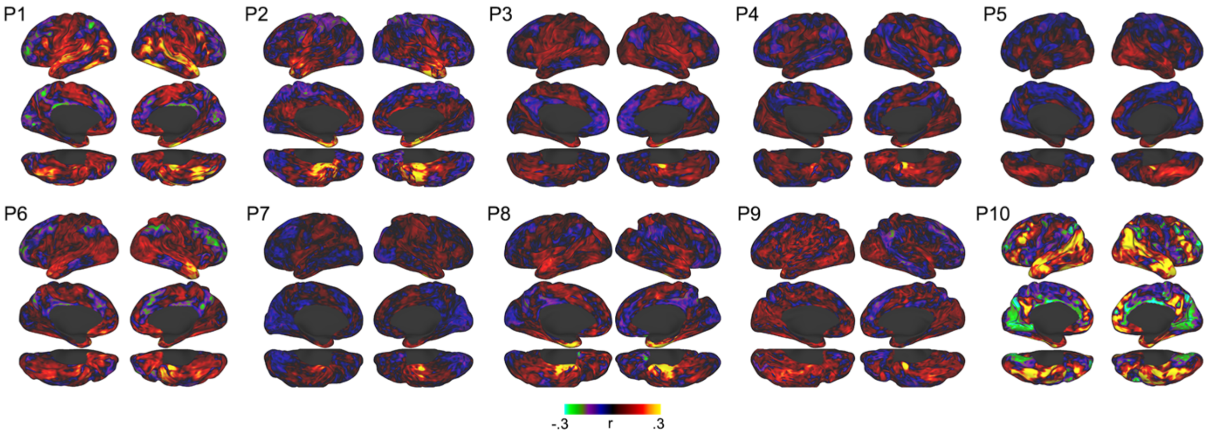

**Figure S8.** Whole-brain resting-state correlation maps from a face-preferring right perirhinal cortex seed region, across all participants.
